# *Sympetrum sanguineum* Newman, 1833 (Odonata, Anisoptera) - only one species with dimorphic females or two sibling species with identical males?

**DOI:** 10.1101/2022.09.17.508356

**Authors:** Babalean Anda Felicia

**Affiliations:** University of Craiova, Faculty of Horticulture, Department of Biology and Environmental Engineering

## Abstract

In this paper it is described a new Sympetrum species - *Sympetrum bigeminus* sp. nov. from Romania. The male of this new species is similar almost to identity with *Sympetrum sanguineum* from which differs only in the more conspicuous lateral thoracic sutures. The females are different from *Sympetrum sanguineum* by the vulvar scale, which is prominent (well visible in lateral view), incurved, with a deep concavity, and shortly bilobed, thus showing morphological characters of a different and distinct species.

## Introduction

Sympetrum Newman, 1833 is a genus of Odonata Anisoptera consisting of around 60 species with a wide geographical distribution, except Australia (Askew, 2004; Wildermuth & Martens, 2019). In Europe, Sympetrum is represented by 11 – 12 species (Askew, 2004; Wildermuth & Martens, 2019) from which 9 species are cited in the Romanian fauna (Cîrdei & Bulimar, 1965): *Sympetrum vulgatum* (Linnaeus, 1758), *Sympetrum flaveolum* (Linnaeus, 1758), *Sympetrum sanguineum* (Müller, 1764), *Sympetrum pedemontanum* (Müller in Allioni, 1766), *Sympetrum danae* (Sulzer, 1776), *Sympetrum fonscolombii* (Selys, 1840), *Sympetrum striolatum* (Charpentier, 1840), *Sympetrum depressiusculum* (Selys, 1841), *Sympetrum meridionale* (Selys, 1841).

The identification keys for species use a set of morphological characters such: pterography, the colour of the legs, anal appendages, male hamuli, female vulvar scale, … The colour of the legs separates the European Sympetrum species in two groups:

a. the black legs group: *S. danae, S. depressiusculum* and *S. sanguineum*
b. the black-yellow stripped legs group: the rest of the species

This paper presents two local populations of black legged Sympetrum, populations in which the males have almost all the morphological characters *of S. sanguineum,* near to identity, while the females have the morphological characters of a different and distinct species. This situation imposes the title question. For the answer to this question, I propose the description of a new species: *Sympetrum bigeminus*.

## Material and method

A total number of 15 individuals with a sex ratio of 3 ♀♀: 12♂♂ were collected in ethanol 80° from 2 populations at hundreds of km distance:

a. Jimbolia – Timiş district: 2 ♀♀, 1 ♂ from a completely dry irrigation canal (06 August 2022) – Fig. 1 and 4♂♂ from the vegetation of a small pool near the Jimbolia recreational swimming pool (07 August 2022). The habitat of the irrigation canal is shared with another Sympetrum species, most probably *S. meridionale*.
b. Craiova – Dolj district: 5♂♂ from the vegetation of a very slow flowing water body in Parcul Romanescu (18 August 2022) and 2♂♂, 1♀ (07 September 2022) from the same location – Fig. 1, where the new species is not associated with another Sympetrum species.

**Fig. 1.**
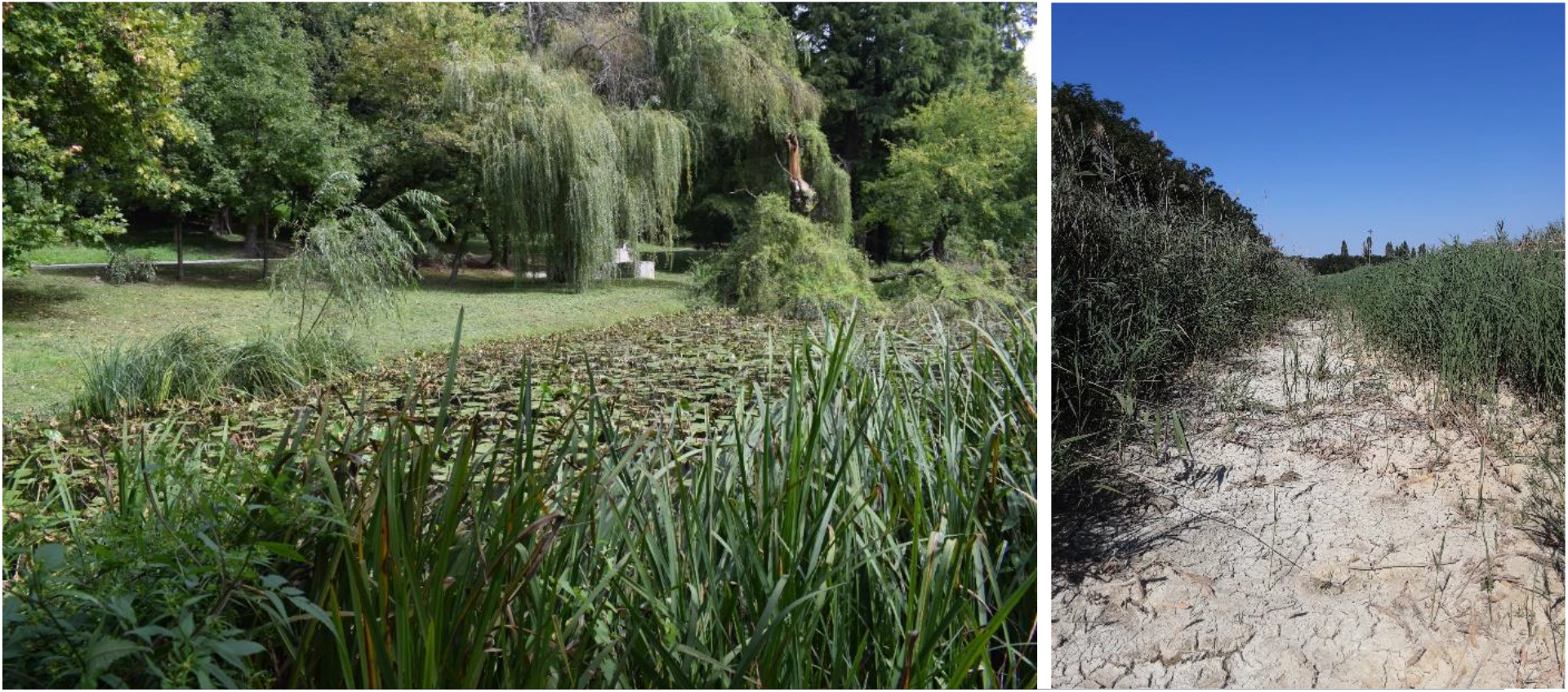
Collecting sites: left – Romanescu Park Craiova, right – dried irrigation canal Jimbolia

The specimens are deposited in author private collection at University of Craiova and will be shared (donated) between Grigore Antipa Museum, Bucharest and elsewhere.

## Results

Systematic

Ord. Odonata Fabricius, 1792

Subord. Anisoptera Selys, 1840

Fam. Libellulidae Selys, 1850

Genus Sympetrum Newman, 1833

### *Sympetrum bigeminus* sp. nov. Babalean

<u>Material examined</u> – type specimens and other specimens Type specimens: *Holotype:* female imago Sbg-H-Jb-Ro-AFB, 06 August 2022, irrigation canal *Paratype no. 1:* female imago Sbg-P1-Jb-Ro-AFB, 06 August 2022, irrigation canal *Paratype no. 2:* female imago Sbg-P2-Cv-Ro-AFB, 07 September 2022 *Paratype no. 3:* male imago Sbg-P3-Jb-Ro-AFB 06 August 2022, irrigation canal *Paratype no. 4:* male imago Sbg-P-Cv-Ro-AFB, 18 August 2022 Where: Sbg – *Sympetrum bigeminus*, H – holotype, P1 – P4 - paratype 1 to paratype 4, Jb – Jimbolia, Cv – Craiova, Ro – Romania, AFB – Anda Felicia Babalean Other specimens – the remaining 11 specimens with the remarque that the identification and assignment of the 4 males from Jimbolia pool to *S. bigeminus* might be uncertain in the absence of the females.
<u>General description</u> General colour of males in living (flight, perching) – intense reddish-brown with an orange-red abdomen – Fig. 2; reddish-brown transparent wings. General colour of females in living – reddish-brown (in two females) and yellow brown (in one female). The smallest male specimen – 31 mm, the largest male specimen – 37 mm.
<u>Diagnosis</u> Male thorax reddish-brown on dorsal and sides, with well-marked black sutures. Dorsal thorax with a broad black anterior band in males and females, producing with the thin black stripe between the two mesepisternum a complete or incomplete T sign. The black ante-ocular stripe present in males and females. A reduced amber patch on forewing and hindwing, better marked on the hindwing. Vulvar scale prominent, incurved, bilobed (Figs. 3.2, 3.10, 3.11-holotype; 4.1, 4.2 – paratype 1; 5.1, 5.2 – paratype 2) with a deep concavity (visible in paratype 1 – Fig. 4.1, 4.2).
<u>Etymology</u> The name of this new species alludes to the male similarity near to identity with *S. sanguineum* (the twin brother).
<u>Description of the female holotype</u> (preserved in ethanol) – Fig. 3 Measurements: total – 33mm

> Head – Figs. 3.4, 3.5, 3.6 Frons yellow with brown, short hairs. Postclypeus, anteclypeus, labrum yellow with brown, short hairs. Mandibles black. Labium yellow and broad. Vertex yellow-brown, hairy. Occiput reddish-brown. The black ante-ocular band (“the moustache” in Smallshire & Swash, 2020) extended to postclypeus, like in *S. vulgatum*. Antenna black. Eyes reddish-brown in the third superior (upper) part and yellow in the third inferior (lower) part.

> Thorax – Fig. 3.1, 3.4, 3.7 Reddish on dorsal. Prothorax with a prominent lobe with very long white setae, typical for Sympetrum. Synthorax: dorsal reddish-brown with a black and large anterior band, lateral yellowish brown with conspicuous black sutures, ventral as in Fig. 3.7.

> Legs – Figs. 3.1, 3.8 Trochanters yellow with a black anterior sclerite of a knee-cap aspect. The second and third pairs of femurs black, the first pair black with a large yellow longitudinal stripe on the front half ventral side.

> Wings – Fig. 3.1, 3.3, 3.9 Reddish-brown in flight, transparent and glittering in the sun light. Forewing and hindwing with a small yellow saffron area, strongly marked on the hindwing. Brown venation. Pterostigma extending in part on two cells; brown in flight, yellow with granular brown dots and with black upper and lower contours under binocular. Cells between radial supplementary vein and posterior margin of both forewing and hindwing 4 – 5 in each oblique series, typical for *S. sanguineum*, as described in Askew (2004) and Boudot et al. (2019).

> Abdomen – Figs. 3.1, 3.2, 3.3, 3.12 Yellow brown with a reddish tint; almost cylindrical, not depressed dorso-ventrally. Without lateral constriction, not club shaped. Two large black spots mid-dorsally on S1 and S2, occupying the breadth of the tergites – Fig. 3.1. Two large mid-dorsally spots on S8 and S9 – Figs. 3.3, 3.12. One large longitudinal black band on each lateral side of segments S3 to S8 – Fig. 3.2. The ninth sternite with long perpendicular hairs, like *S. sanguineum* – Fig. 3.10.

> Vulvar scale – Fig. 3.2, 3.10, 3.11 Prominent, very well visible in lateral view. Incurved – curved internally towards the sternites, with a deep concavity. Consists of two plates – a smaller external triangular plate and a larger internal, shortly bilobed plate, the lobes triangular.

> Anal appendages – Fig. 3.3, 3.12 Broader than those figured by Cîrdei & Bulimar (1965).

**Fig. 2.**
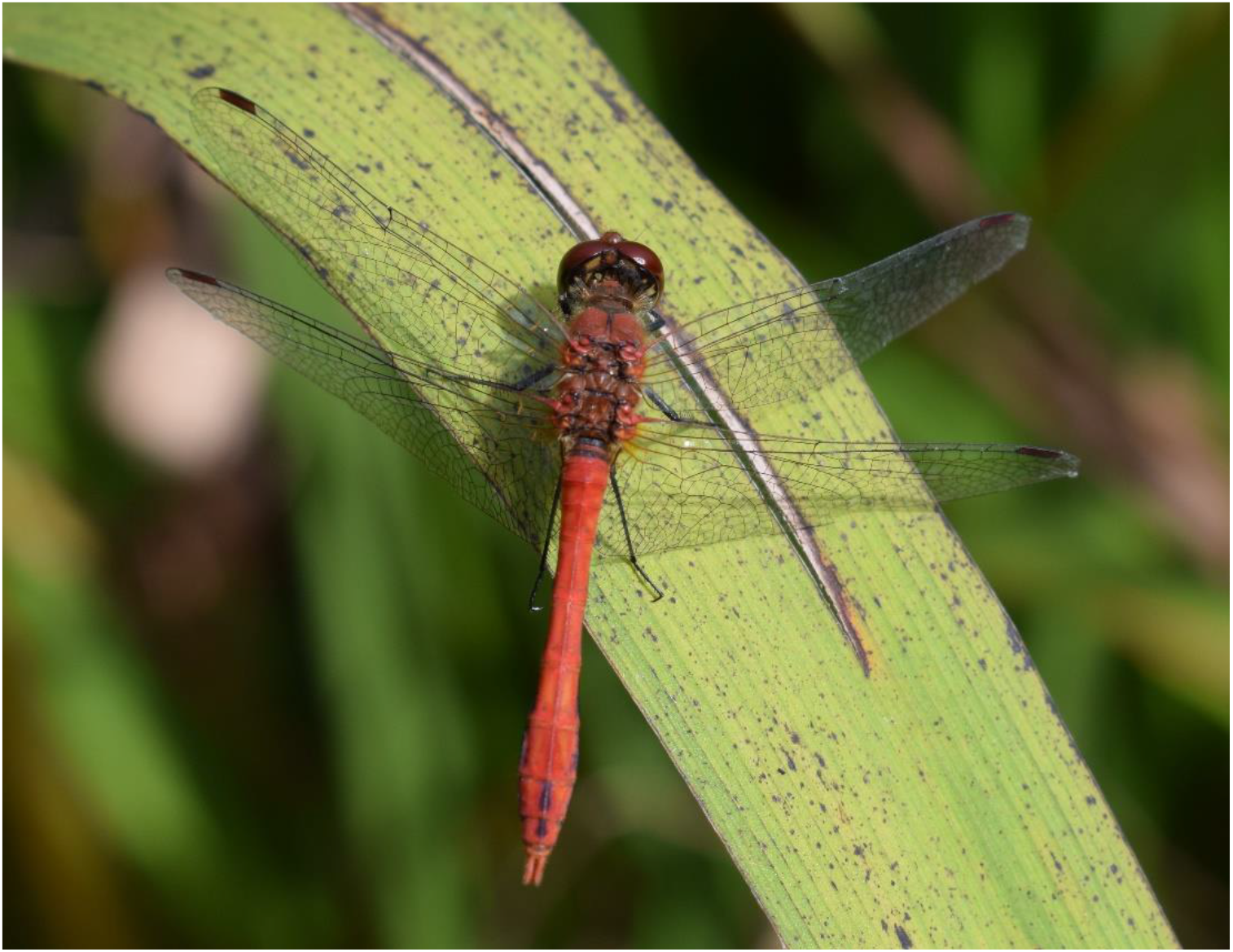
General habitus of a male *Sympetrum bigeminus* (photo by Anda Felicia Babalean on 15 September 2022)

**Fig. 3.1.**
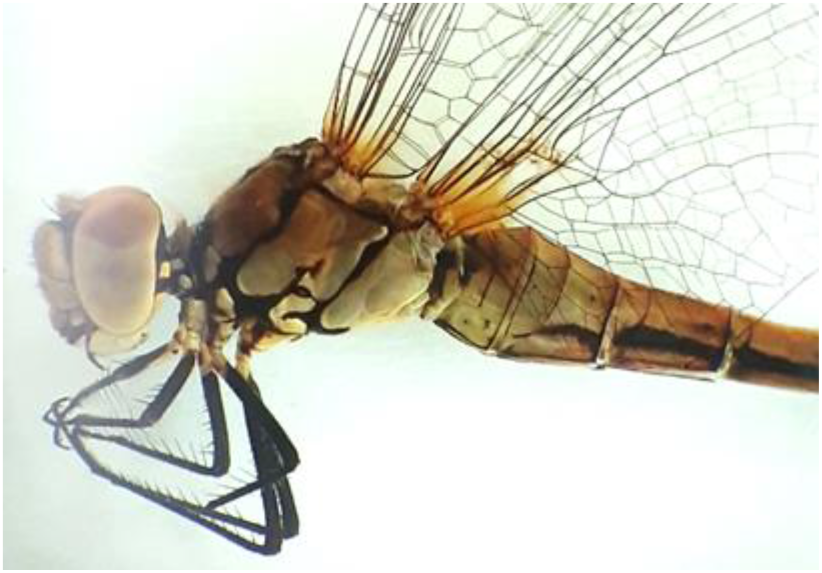
*Sympetrum bigeminus* – holotype Head, thorax, legs, wings, abdomen

**Fig. 3.2.**
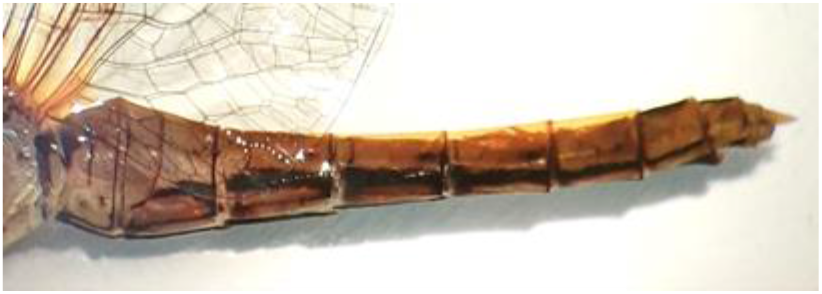
*Sympetrum bigeminus* – holotype Abdomen in lateral view

**Fig. 3.3.**
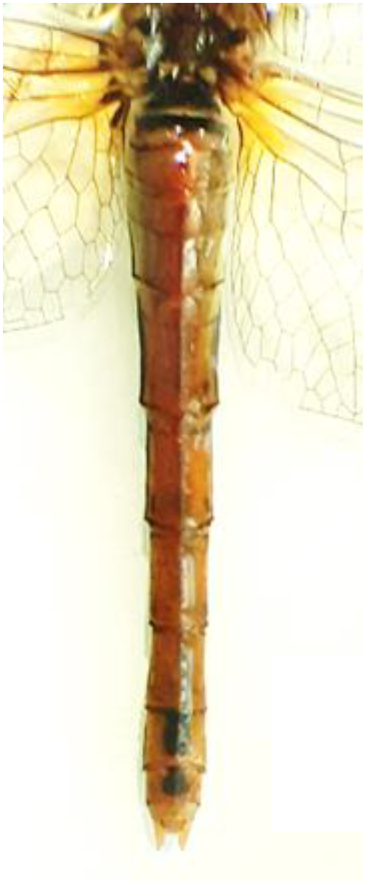
*Sympetrum bigeminus* – holotype Abdomen in dorsal view

**Fig. 3.4.**
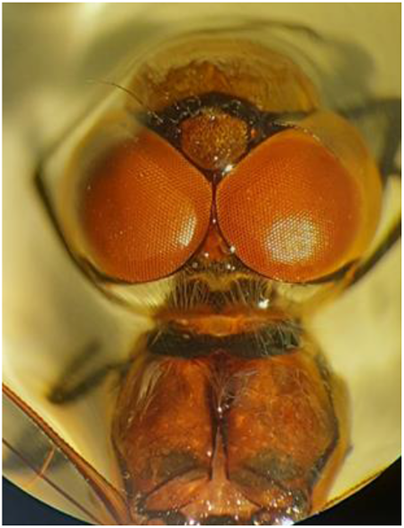
*Sympetrum bigeminus* – holotype Head and thorax in dorsal view

**Fig. 3.5.**
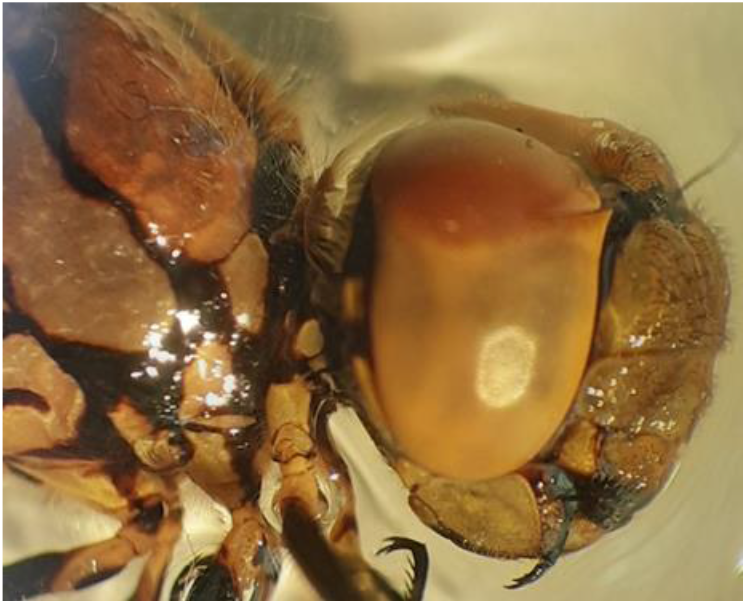
*Sympetrum bigeminus* – holotype Head in lateral view

**Fig. 3.6.**
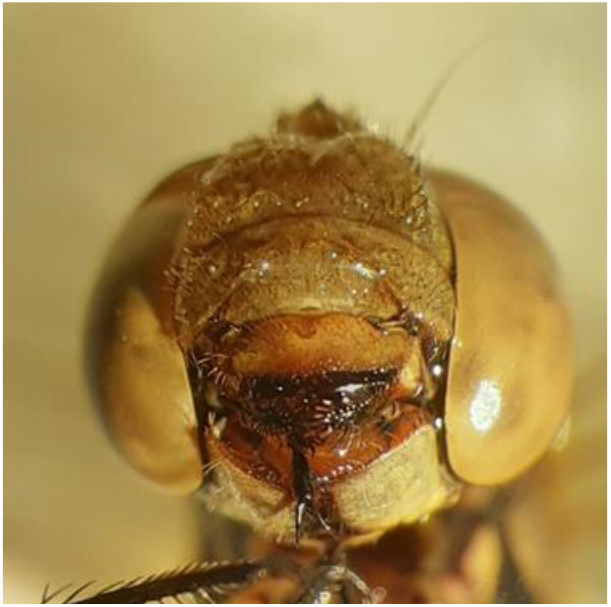
*Sympetrum bigeminus* – holotype Head in frontal view

**Fig. 3.7.**
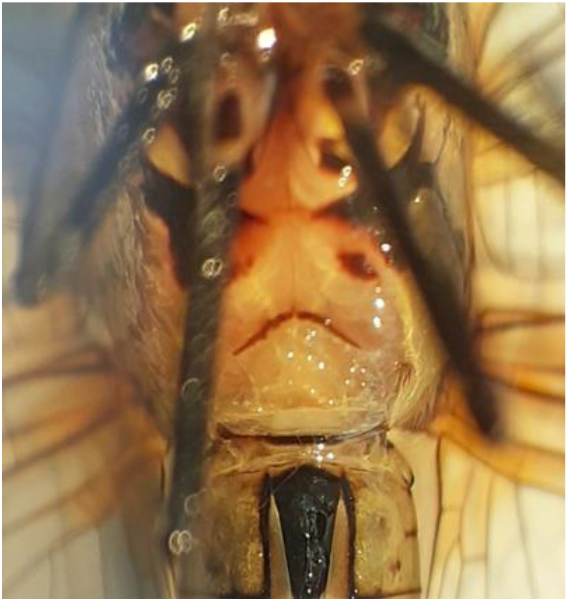
*Sympetrum bigeminus* – holotype Thorax in ventral view

**Fig. 3.8.**
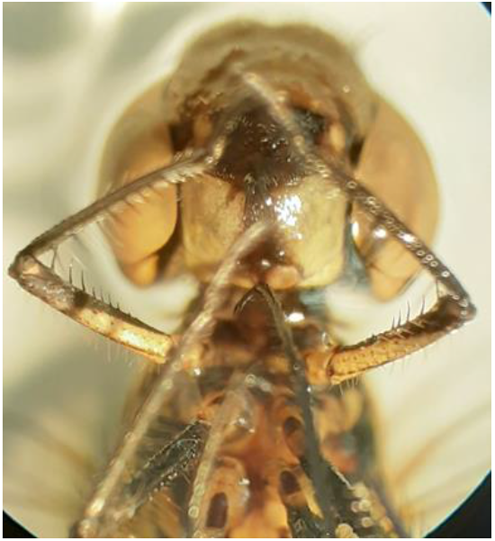
*Sympetrum bigeminus* – holotype Front femora in ventral view

**Fig. 3.9.**
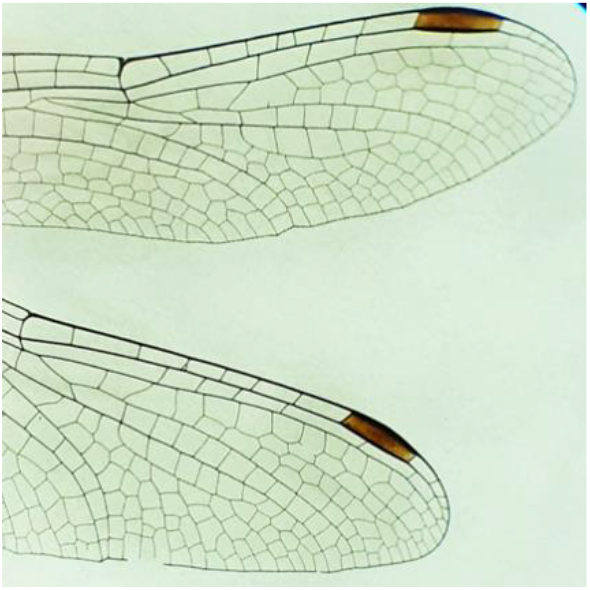
*Sympetrum bigeminus* – holotype Wings

**Fig. 3.10.**
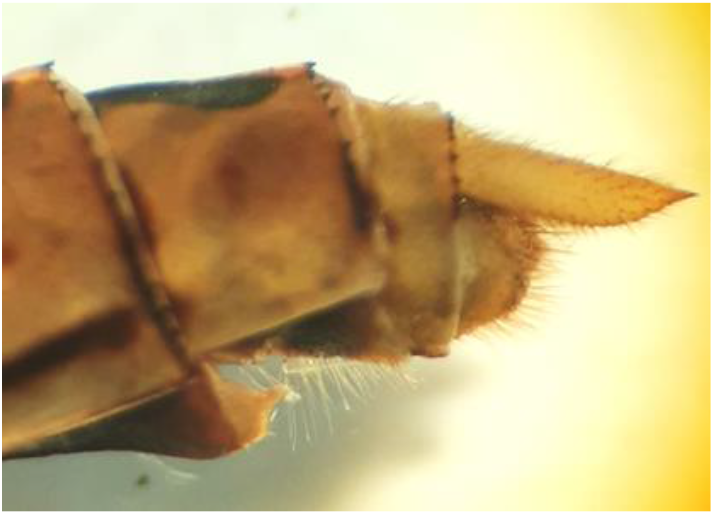
*Sympetrum bigeminus* – holotype Vulvar scale in lateral view

**Fig. 3.11.**
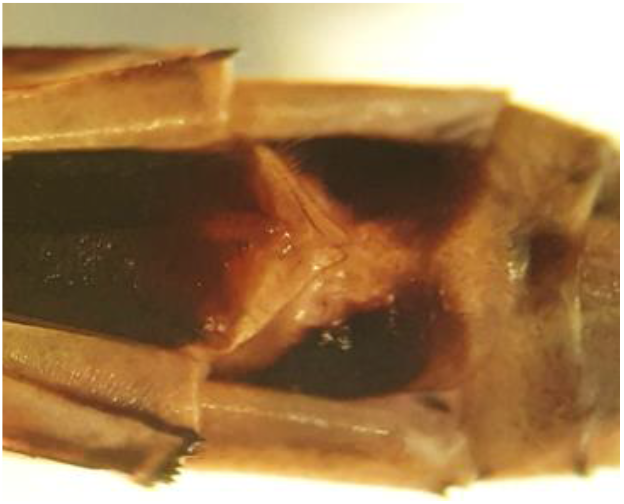
*Sympetrum bigeminus* – holotype Vulvar scale in ventral view

**Fig. 3.12.**
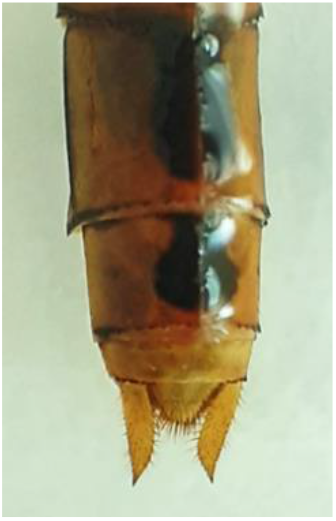
*Sympetrum bigeminus* – holotype Anal appendages in dorsal view

**Fig. 4.1.**
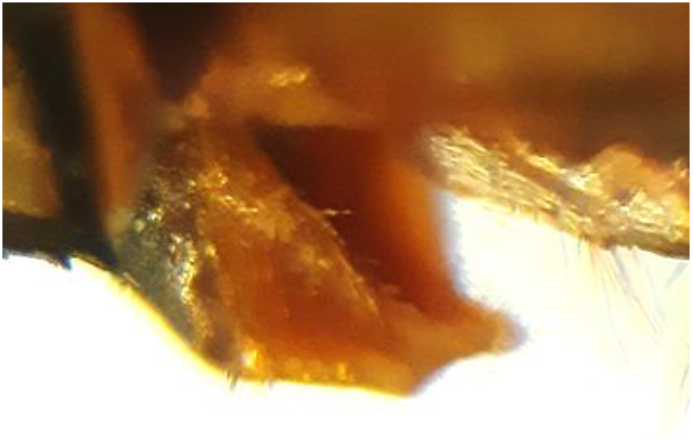
*Sympetrum bigeminus* – paratype 1 Vulvar scale in lateral view showing the concavity

**Fig. 4.2.**
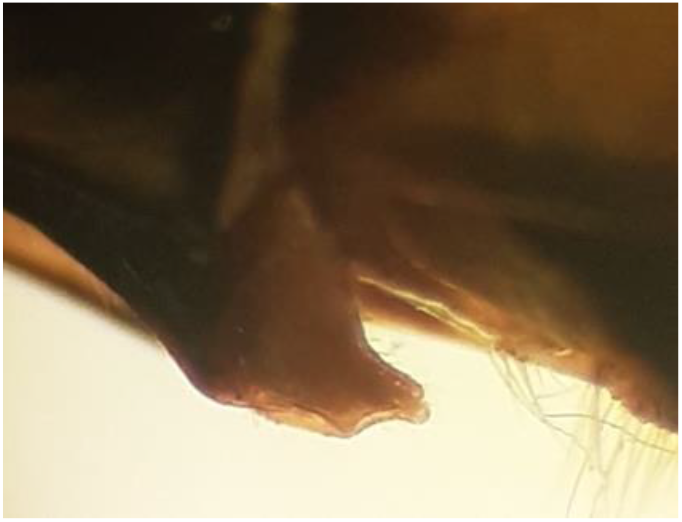
*Sympetrum bigeminus* – paratype 1 Vulvar scale in lateral view showing the 2 lobes

**Fig. 5.1.**
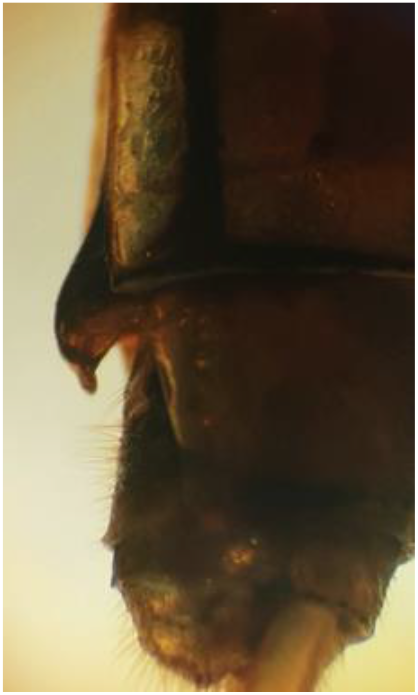
*Sympetrum bigeminus* – paratype 2 Vulvar scale in lateral view

**Fig. 5.2.**
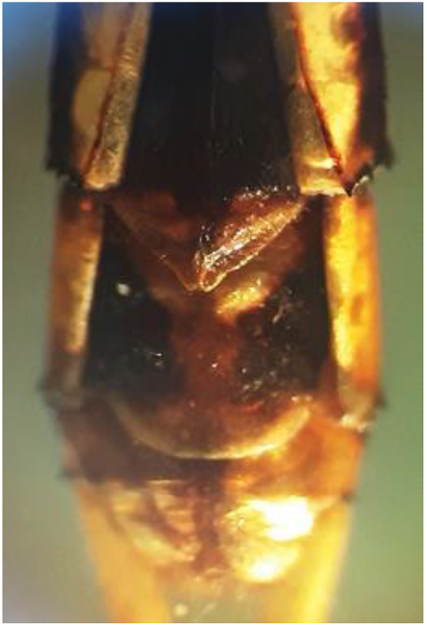
*Sympetrum bigeminus* – paratype 2 Vulvar scale in ventral view showing the 2 lobes

<u>Description of the male paratype 3 (preserved in ethanol) – Fig. 6</u> Measurements: total – 35 mm.

> Head – Figs. 6.1, 6.4, 6.5 All the components of the head are reddish-brown, except the black mandibles. Frons intense red in living specimens. The black ante-ocular band extended to postclypeus. Antenna black. Eyes reddish-brown with a much redder tint in the third superior part.

> Thorax – Fig. 6.1 Synthorax: reddish-brown, more intense red tint on dorsal, with a black anterior large band and with conspicuous black lateral sutures.

> Legs – femora and tibiae entirely black.
>
> Wings – Figs. 6.6, 6.7 Very similar with those of the females.

> Abdomen – Figs. 6.2, 6.3 Reddish-brown and club shaped, with a marked constriction at S4. Two large black spots mid-dorsally on S1 and S2. The general aspect is identical with the abdomen of *S. sanguineum* as illustrated in literature (Boudot et al., 2019; Wildermuth & Martens, 2019; Smallshire & Swash, 2020).

> Accessory genitalia (hamuli) – Fig. 6.8 The two hamular processes are of about equal length, the inner anterior process has a curved black tip. The aedeagus is the flagellar type and is composed by an internal impair black “*flagellum*” and two white lateral “*flagella*” – Fig. 6.8.

**Fig. 6.1.**
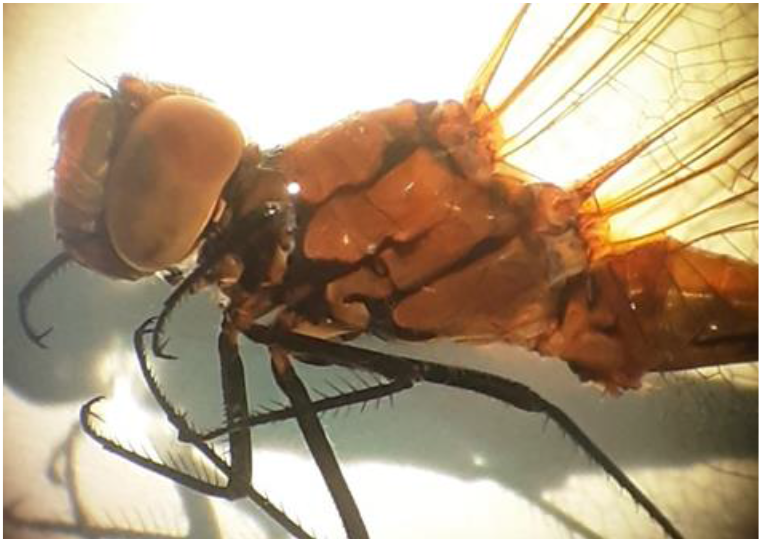
*Sympetrum bigeminus* – paratype 3 Head, thorax, legs in lateral view

**Fig. 6.2.**
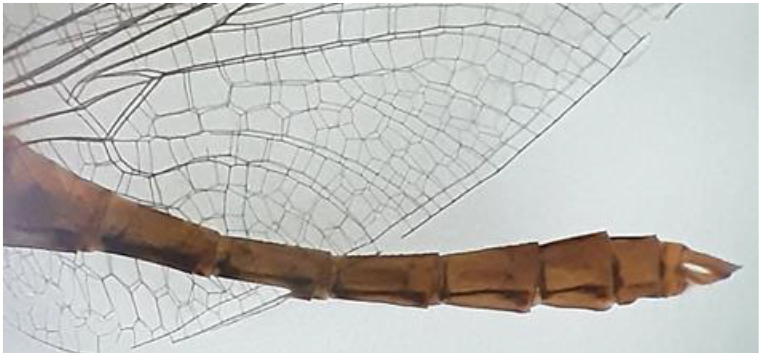
*Sympetrum bigeminus* – paratype 3 Abdomen in lateral view

**Fig. 6.3.**
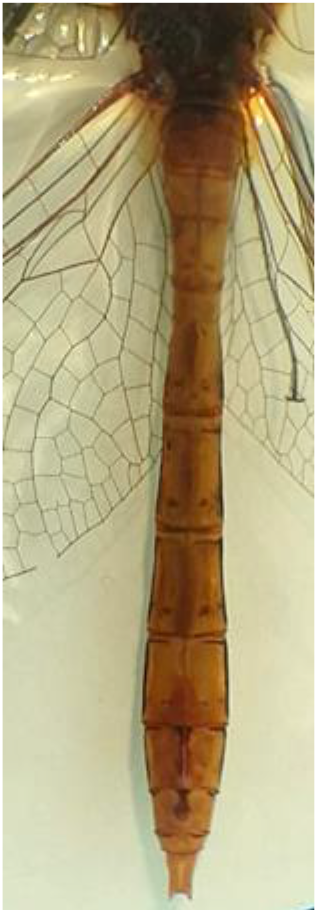
*Sympetrum bigeminus* – paratype 3 Abdomen in dorsal view

**Fig. 6.4.**
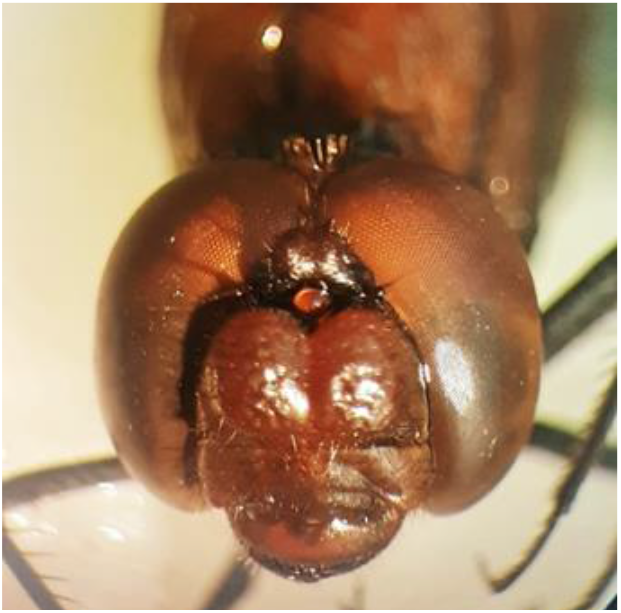
*Sympetrum bigeminus* – paratype 3 Head in frontal view

**Fig. 6.5.**
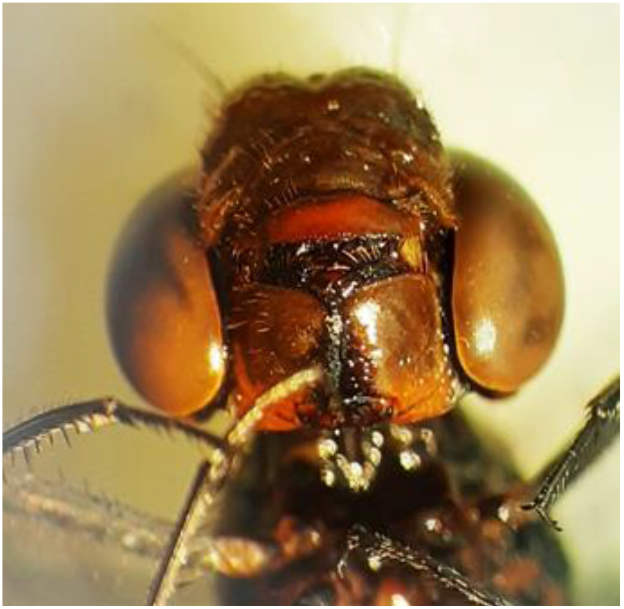
*Sympetrum bigeminus* – paratype 3 Head in ventral view

**Fig. 6.6.**
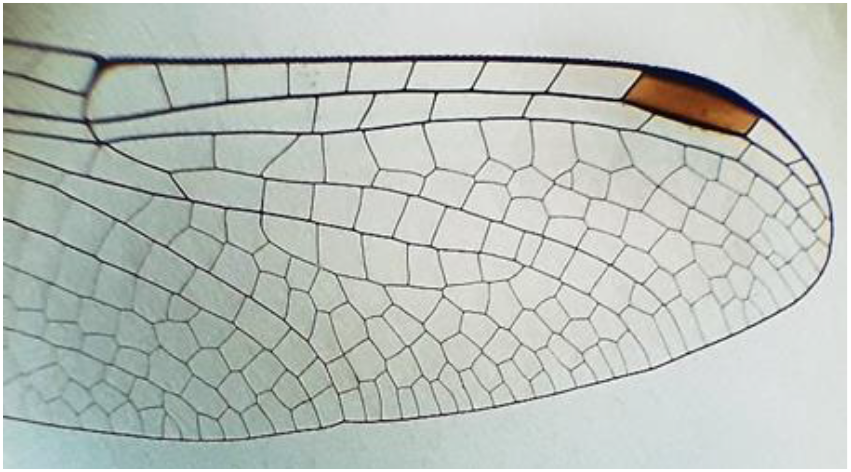
*Sympetrum bigeminus* – paratype 3 Forewing

**Fig. 6.7.**
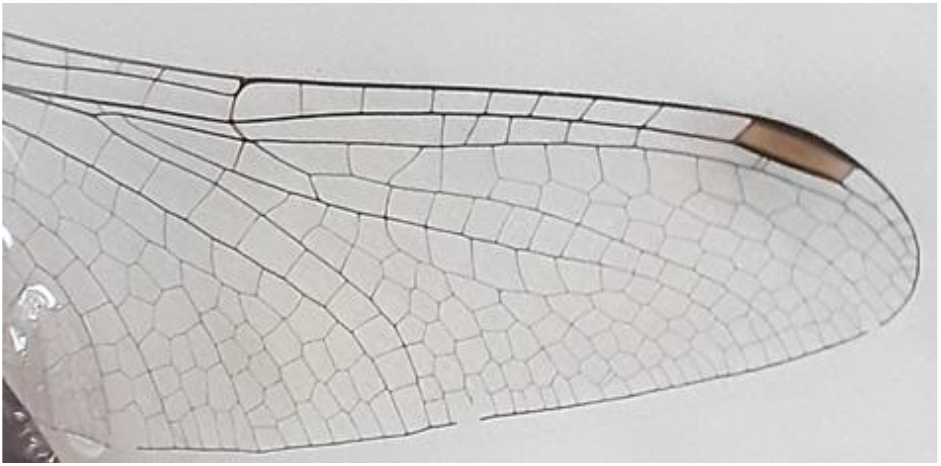
*Sympetrum bigeminus* – paratype 3 Hindwing

**Fig. 6.8.**
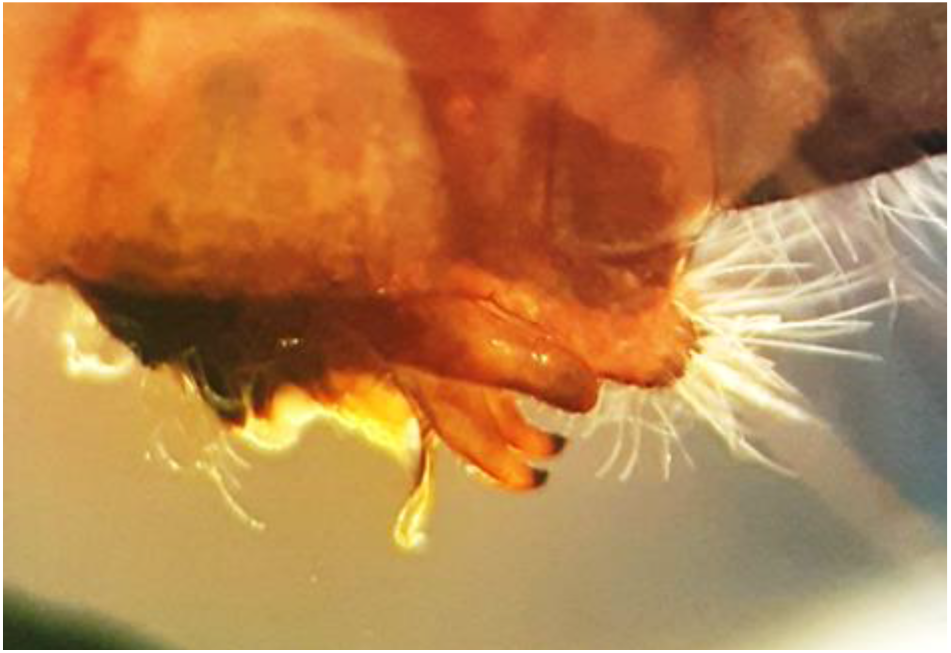
*Sympetrum bigeminus* – paratype 3 Hamular processes, ligula and aedeagus in lateral view

The variability of the male abdomen – Fig. 7

> The shape (club-shape) is a constant character in both populations (Jimbolia and Craiova).
>
> The pattern of the black spots on lateral abdomen is a variable, individual character, with spots of various size to no spot – Fig. 7.1, 7.2, 7.3.

**Fig. 7.1.**
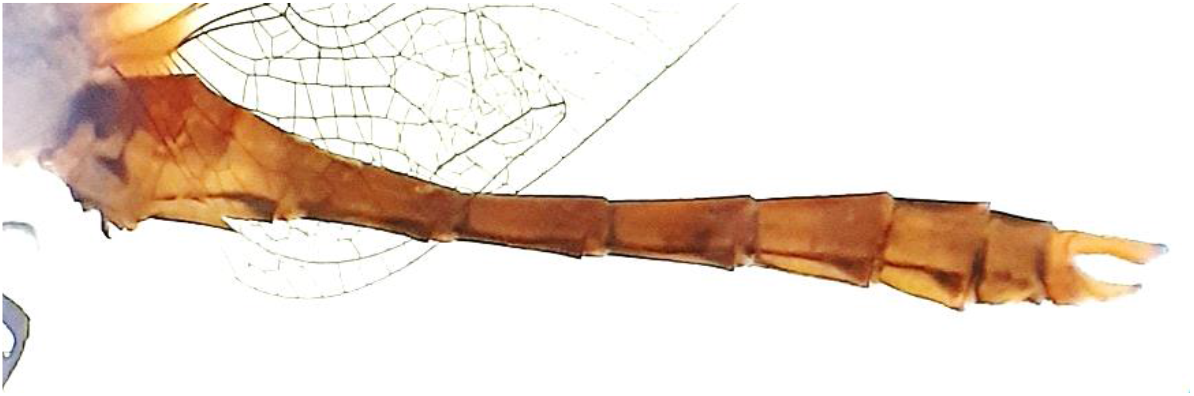
*Sympetrum bigeminus* – male from Craiova population Abdomen in lateral view in lateral view

**Fig. 7.2.**
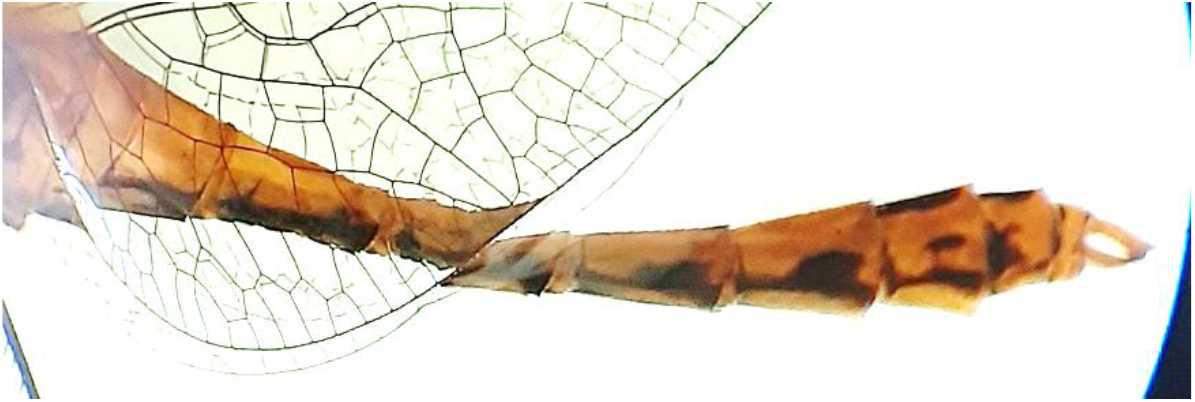
*Sympetrum bigeminus* – male from Jimbolia population Abdomen in lateral view in lateral view

**Fig. 7.3.**
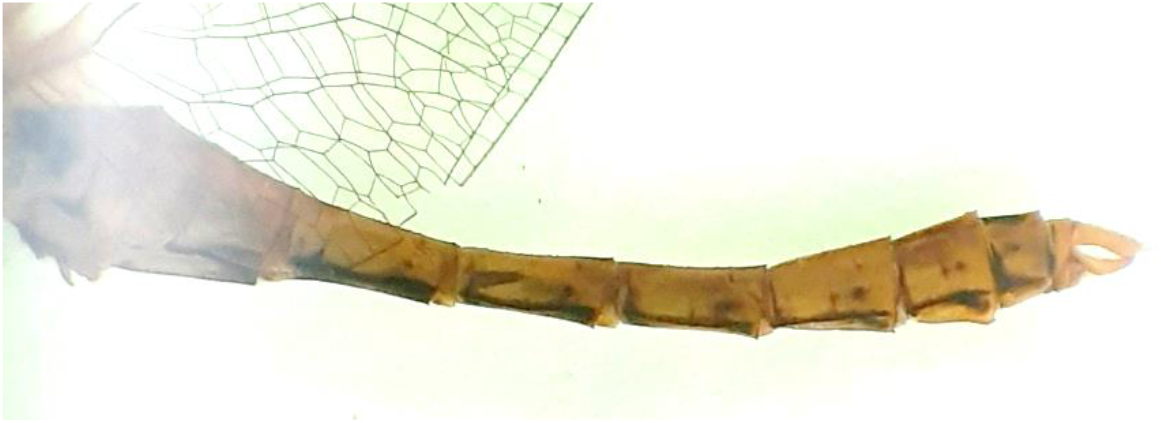
*Sympetrum bigeminus* – male from Jimbolia population Abdomen in lateral view in lateral view

The variability of the accessory genitalia – Figs. 8, 9

> The structure of the hamuli is a constant character in both populations (Jimbolia and Craiova): the two hamular processes are of about equal length, the inner anterior process with a curved pointed black tip – Fig. 8, 9.
>
> The shape of the outer posterior process is a variable, individual character.
>
> The position of the ligula is a variable character between the two populations. Compared with the literature (Askew, 2004, Fig. 325, pg. 174), the ligula is oriented in a *S. danae* fashion in the population from Jimbolia – Fig. 8.

**Fig. 8.**
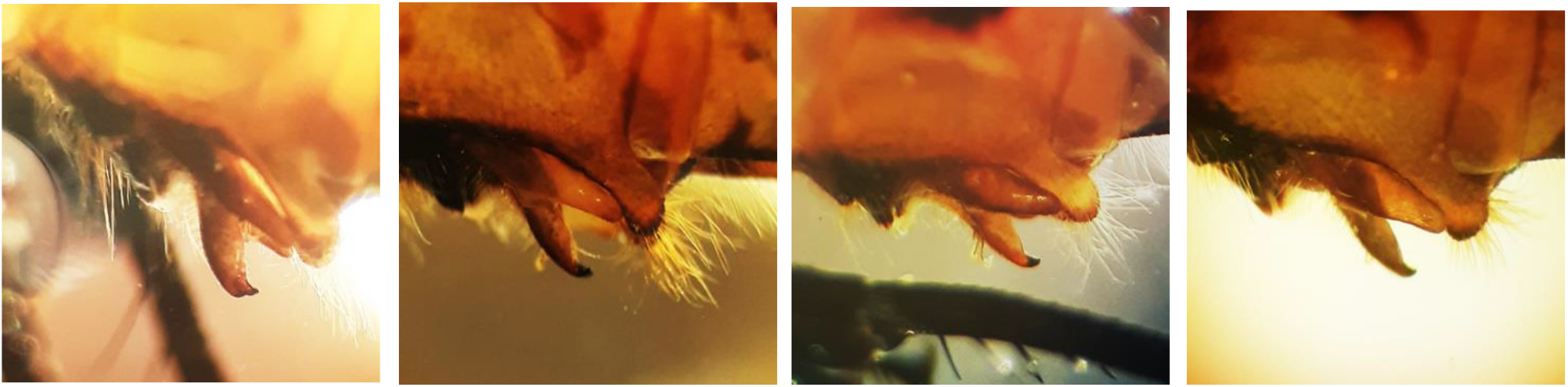
*Sympetrum bigeminus* – the variability of accessory genitalia in 4 specimens from Jimbolia population

**Fig. 9.**
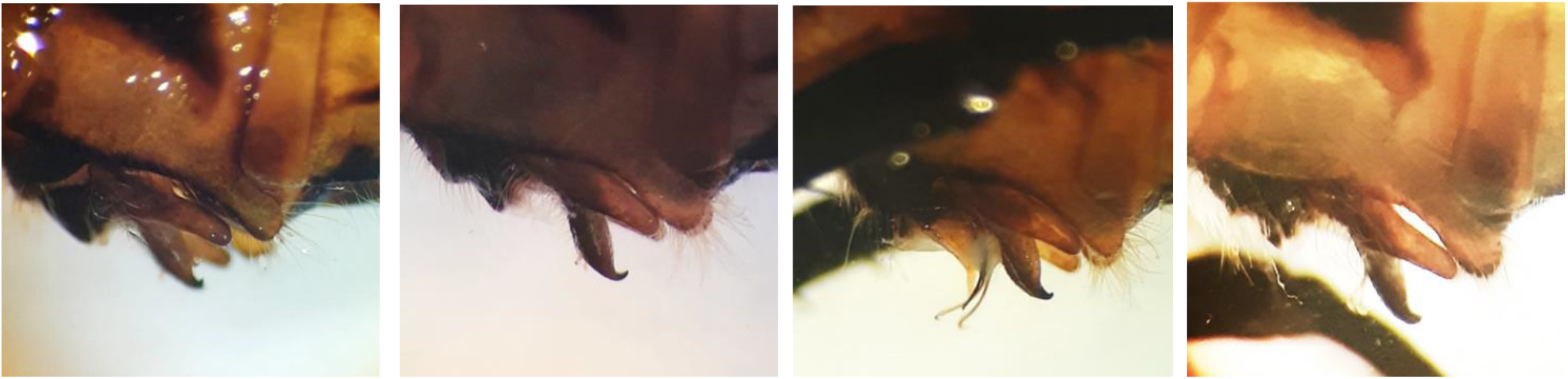
*Sympetrum bigeminus* – the variability of accessory genitalia in 4 specimens from Craiova population

## Discussions

The colour of the body may be influenced by some factors like the light (sun, binocular bulb), solubility of some pigments in ethanol, depigmentation in time, etc. Thus, the perception and description of the colour in text and illustration may not be neither of perfect accuracy nor consistent (for all researchers). For instance, the thorax of *S. sanguineum* is described with yellow sides (Askew, 2004) or yellow brown sides (Boudot et al., 2019) and with black markings and is illustrated as intense orange-red in Wildermuth & Martens (2019) and in Smallshire & Swash (2020). Nevertheless, the male specimens from Craiova population carefully observed in living (flight and perching) were entirely reddish brown with the abdomen orange-red and frons deep red.

The only character that distinguishes the males of *Sympetrum bigeminus* from *Sympetrum sanguineum* is the more conspicuous black sutural pattern on sides of thorax.

The possibility that the males and the females described as *S. bigeminus* belong to different species sharing the same habitat (males described as *S. bigeminus* would be in fact *S. sanguineum*; females described as *S. bigeminus* would belong to a different known species) is annulled by the following:

- the prominent and bilobed vulvar scale of *S. bigeminus* is very different from that of *S. sanguineum* which “is neither prominent, nor bilobed” (Askew, 2004). The bilobed vulvar scale of *S. bigeminus* is related with *S. flaveolum* and *S. pedemontanum*. *S. bigeminus* differs from both *S. flaveolum* and *S. pedemontanum: S. flaveolum* is a species illustrated with stripped, black-yellow legs (Wildermuth & Martens, 2019); *S. pedemontanum* is an unmistakable species by the broad reddish-brown transverse band on the wing.
- males and females in Craiova population are found together and do not mix with another Sympetrum species – no specimen of another Sympetrum species was collected from Craiova so far.
- the females described as *S. bigeminus* are associated with males described as *S. bigeminus* in both Jimbolia (irrigation canal) and Craiova populations. Nevertheless, in the absence of the collected females, the 4 male specimens from the Jimbolia pool might be attributed as well to *S. sanguineum*.

